# A Conditional Random Field approach for de novo reconstruction of bacterial haplotypes from a de Bruijn graph representation

**DOI:** 10.64898/2026.05.11.724222

**Authors:** Aranka Steyaert, Marie Van Hecke, Kathleen Marchal, Jan Fostier

## Abstract

**Background:** Detecting distinct bacterial strains in a mixed sample is an important, yet less well-developed aspect of metagenomic research. Several methods exist that successfully retrieve a de novo reconstruction of viral strains. However, the reconstruction of bacterial haplotypes poses its own distinct challenges, and methods that successfully reconstruct full genome-length bacterial strains de novo are scarce. Here, we develop HaploDetox, a method for de novo bacterial haplotype reconstruction from short reads. We use a de Bruijn graph representation of the reads in which nodes correspond with k-mers from the read set and arcs represent overlap between two nodes’ sequences. Our aim is to accurately assign labels to each node and arc in the graph to reveal the presence or absence of their corresponding sequence in individual strains.

**Results:** Using a negative binomial mixture model, we model the relationship between the read coverage of nodes and arcs in the graph and their presence in a strain. We achieve improved labelling accuracy by including contextual information from neighbouring nodes and arcs with a Conditional Random Field. These labels are used to extract strain-specific de Bruijn graphs from the original graph. Additionally, we allow users to assess the number of strains present in the dataset based on model selection criteria. We evaluate our node/arc labelling accuracy on simulated datasets and in silico mixes of real datasets containing different numbers of strains, as well as on in vitro mixed real datasets. Existing de novo haplotype reconstruction methods present their reconstruction as strain-specific sets of SNPs. We demonstrate that HaploDetox assigns strain-specific SNPs with a higher recall and similar precision than existing methods, by aligning the unitigs from strain-specific graphs to a reference genome.

**Conclusions:** We achieve improved strain-specific SNP phasing accuracy as compared to existing methods for de novo bacterial haplotype reconstruction. Additionally, HaploDetox is not limited to the determination of strain-specific SNPs, and other types of variant calls can be obtained through reference alignment. Finally, strain-specific de Bruijn graphs are an important first step towards full genome-length bacterial haplotype-aware assembly.

## Background

Recent technological advances enabled the generation of whole genome sequencing data to study metagenomic microbial samples or clonal bacterial populations. Second generation short Illumina sequencing reads still make up the majority of data used for such analyses [1]. Proper identification of within-species variation is important to study mixed infections, antibiotic resistance, and evolutionary or spatial dynamics of a species [2]. The precise definition of a bacterial or microbial strain has been extensively discussed in literature, see e.g. [3, 4]. Here, the term strain refers to a member of a collection of genomes with sub-species level variation. Determining the genomic sequence of each of these co-existing strains in a sample will be referred to as *bacterial haplotype reconstruction* or haplotype-aware assembly. While many methods exist to determine and/or assemble different species within a metagenomic sample, methods that discern the presence of multiple bacterial strains are still in their infancy [5].

Here, we focus on *de novo* whole genome reconstruction of strains of a single species in a bacterial dataset, without using a reference genome. State of the art de novo haplotype reconstruction methods based on short reads such as EVORhA [6], BHap [7], and mixtureS [8] still use a reference genome to define haplotypes based on variants called with respect to this reference. They are considered de novo methods because they do not require a database of individual strains. However, BHap and mixtureS only determine the SNPs present in the strains and ignore other types of mutations, thus not characterising the observed strains fully. EVORhA does include all types of mutations, but was evaluated as the least accurate of these three methods in a recent review by Ventolero et al. [5]. Other approaches to bacterial haplotype reconstruction exist; we refer to [2] and [5] for a comprehensive discussion. Many of these approaches rely on a reference database of strain genomes against which reads of a mixed sample are aligned. Alternatively, many existing de novo techniques only identify variation in marker genes or other conserved regions.

While de novo bacterial haplotype reconstruction still faces many challenges, viral strains can often be successfully reconstructed from mixed sample sequencing data, even without the use of a reference genome [9]. However, methods aimed at viral data do not scale well for mixed bacterial strains, which are larger and more repeat-rich than viruses [10]. Moreover, properties of viral genomes that are exploited by viral haplotype reconstruction tools, such as a high divergence rate between sequences, do not hold for bacterial strains [6]. Because of the high mutation rate in viruses, even short reads can be informative to assign mutations to their respective strain. In contrast, the low mutation rate in bacteria renders the phasing of strains based on co-occurring variants less effective [5].

Here, we develop a method that can discern the full genome sequences of strains of the same species present in a bacterial short read dataset. When starting from a metagenomic dataset, an appropriate (sub)set of reads can be obtained by binning the reads at species level. For a fully de novo approach, bins can be created by mapping the reads to Metagenome Assembled Genomes (MAGs) obtained from a metagenomic assembler. We efficiently represent the reads and their overlaps with a de Bruijn graph and we phase this graph directly by determining each node’s and arc’s presence in a strain genome. Based on these strain assignments, we extract *strain-specific de Bruijn graphs* that can be used to call strain-specific variants, but are also an important first step towards full haplotype-aware assemblies.

### Details on related existing methods

EVORhA [6] requires a reference genome and variant calls of the read set to this reference. In a first step, small windows containing potential haplotypes are created. These haplotypes are determined by consecutive polymorphisms that co-occur in reads covering the window. For each window, the read support is determined and polymorphisms with low support are assumed to have arisen from sequencing errors and are not considered further. In the next step, windows that share polymorphisms, i.e. that overlap, are concatenated. Finally, these maximally extended haplotype windows are combined into genome-wide haplotypes based on the frequencies of the haplotypes in a window. Under the assumption that the observed frequencies are generated by a mixture of Gaussians with the true frequencies as mean, haplotypes from different windows with similar frequency are clustered together.

BHap [7] uses a de Bruijn graph to obtain contigs from the read set. A flow network is constructed that represents each contig with a node: the contigs are mapped to a reference genome and two nodes are connected by a directed arc if their contigs map to adjacent regions in the reference genome. The flow capacities are determined by the read coverage of the contigs. A set of haplotype-specific read coverages is determined based on a mixture of Poissons fitted to the capacities of the flow network. The flow network is decomposed by greedily extracting paths such that the read support does not exceed the capacity through the arcs in the path. The read support is subtracted from the arc capacities used by the path and the process is repeated. Each path in the decomposition represents a genome-wide haplotype. This whole procedure is repeated for different *k*-mer sizes and the results are clustered and combined to obtain a single consensus set of haplotypes.

MixtureS [8] maps the read set to a reference genome and identifies all polymorphic positions as well as the frequency with which each variant at a polymorphic position occurs. Only biallelic variants are considered. Polymorphic positions that likely correspond with sequencing errors are filtered based on read coverage criteria. An Expectation-Maximisation algorithm is used to estimate the probability that each polymorphic position’s non-reference allele belongs to a certain strain given an estimated relative abundance of the different strains.

## Methods

Fig. 1 illustrates the concepts introduced here. We assume that our dataset is a collection of reads originating from *S* different strain genomes. Each strain *s* is present in the dataset with an abundance that is a fraction *π*_*s*_ of the total genetic content. Hence, each strain will be sequenced at a sequencing depth that is a fraction *π*_*s*_ of the overall sequencing depth. From this set of reads we construct a de Bruijn graph. All length *k* subsequences of the reads, called *k*-mers, correspond with nodes in the de Bruijn graph. Two nodes are connected by a directed arc when they share a *k*−1 length overlap and the *k* + 1-mer formed by concatenating the first character of the source node, the overlapping sequence, and the last character of the target node is present in the reads. Each *k*-mer resp. *k* +1-mer is present in the reads a certain number of times; we call this number the *coverage* of a node, resp. arc. For ease of notation we will use the term *k*-mer when referring to both nodes and arcs. However, the sequence represented by an arc has one more character than a sequence represented by a node, and our model does take this into account.

**Fig. 1.**
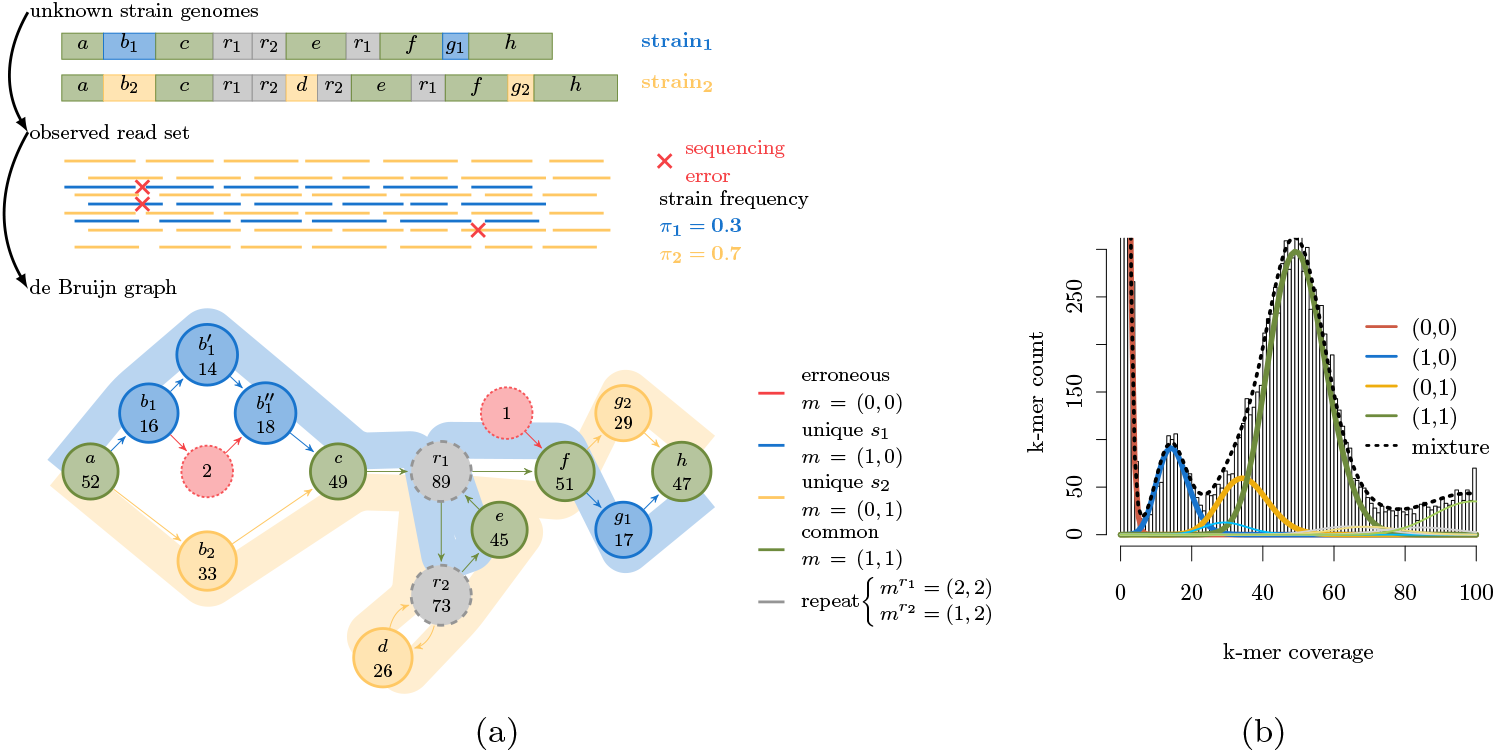
Illustration of the haplotype-aware assembly problem with two strains. Given a set of reads originating from two strain genomes, we reconstruct both genomes using a de Bruijn graph. The *k*mer coverage of the nodes (and *k* + 1-mer coverage of the arcs (not shown)) depends on the frequency with which the strains occur in the sample, the total sequencing depth, and on the multiplicity with which the sequence occurs in the original genome. **(a)** Two (unknown) strain genomes, reads that contain sequencing errors, and the corresponding de Bruijn graph. Sequencing errors result in spurious nodes and arcs in the graph. Sequence *b*_1_ is split into three nodes because of the additional variation introduced by the sequencing errors. The walks that represent the original genomes are shown in blue, resp. yellow in the background of the graph. Nodes and arcs are coloured according to their multiplicity (all repeats are coloured gray regardless of their multiplicity (see legend)). **(b)** *k*-mer coverage histogram with a two strain mixture model of the coverage-multiplicity relationship. The red curve (multiplicity (0, 0) in the legend) captures k-mers with a very low coverage that arise due to sequencing errors with a negative binomial distribution. The other curves represent negative binomial distributions centred around the average coverage of their corresponding multiplicity. Curves corresponding with distributions that represent repeat-multiplicities are drawn thinner and are not in the legend.

The de Bruijn graph is a superposition of multiple strain genomes: each genome is present as a walk through the graph, assuming that there are no coverage gaps (Fig. 1(a)). The *strain-specific multiplicity* denotes the number of times the sequence corresponding with a node/arc is present in the genome of a single strain. The *multiplicity of a node/arc x* is a vector 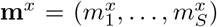, with 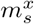 the strain-specific multiplicity of *x* in strain *s*. In the presence of sequencing errors, a node/arc can also have a multiplicity of **m**^*x*^ = (0, …, 0), i.e., when its corresponding sequence is not truly present in any genome from the sample. The observed coverage of a *k*-mer is related to its multiplicity: *k*-mers that occur with a higher multiplicity or in more than one strain will have a higher coverage than sequences unique to a single strain. The *k-mer spectrum* or range of observed *k*-mer coverage values is visualised with a histogram in which *k*-mers with a certain multiplicity can be observed as peaks around a coverage value, see Fig. 1(b). The histogram has an additional peak around *k*-mer coverage ≈ 1 caused by spurious *k*-mers that are present because of sequencing errors. We model this with a mixture of negative binomial distributions, such that each distribution in the mixture corresponds with a multiplicity **m**. Because each strain genome corresponds to a walk in the graph, every node that is entered by an incoming arc must be exited by exactly one outgoing arc. This results in a *conservation of flow of multiplicity property* :

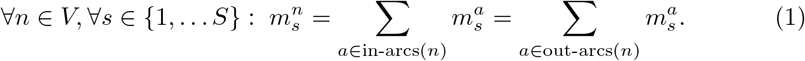

Our aim is to assign strain-specific multiplicities to all nodes and arcs in the de Bruijn graph. We model the strain-specific multiplicity assignment problem with a *Conditional Random Field (CRF)* [11]; such models allow to include local contextual information on how variables interact (here expressed by the conservation of flow property). We encode strain-specific multiplicities and the observed coverage of nodes/arcs as variables, while the coverage-multiplicity relationship and the conservation of flow within a strain (eq. (1)) are encoded as probabilistic relations between variables. Multiplicities are used to determine strain-specific de Bruijn graphs. We call our method HaploDetox. This is an extension of our model of de Bruijn graphs for bacterial isolates in [12, 13], we refer the reader to these works for more information on modelling de Bruijn graphs with CRFs. In the remainder of this section we describe the HaploDetox pipeline, and our evaluation methodology.

### HaploDetox pipeline

HaploDetox consists of several stages (Fig. 2). In stage 1, we construct a de Bruijn graph with BCALM 2 [14] and compute the coverage of all nodes and arcs. In stage 2, we define a CRF based on a mixture model and conservation of flow of multiplicity for a fixed number of strains S. We fit an initial mixture model to the observed *k*-mer spectrum with expectation-maximisation (EM). We infer multiplicity assignments from the CRF by computing marginal probabilities for the multiplicity labels of nodes and arcs. Depending on the size of the CRF, we use exact (variable elimination) or approximate (loopy belief propagation) inference techniques, see also [13]. To reduce the number of nodes and arcs, we use the multiplicity labels from a CRF with the initial mixture model to remove likely erroneous nodes and arcs from the graph. Afterwards, we train the final mixture model using EM with multiplicity labels based on the CRF to improve the model fit. When no estimated number of strains is passed to HaploDetox, models under different strain number assumptions are fitted and the best model is selected using the Bayesian Information Criterion (BIC). Because of the increasing complexity of the CRF for increasing number of strains, we only test *S* ∈ {2, 3, 4} . In stage 3, multiplicities are inferred for all nodes and arcs in the de Bruijn graph. These multiplicities are used to phase the de Bruijn graph into several strain-specific graphs. Below we give more details on the techniques used in stage 2 and 3, further mathematical details for the model training and number of strains selection are provided in Additional file 1, Appendix A.

**Fig. 2.**
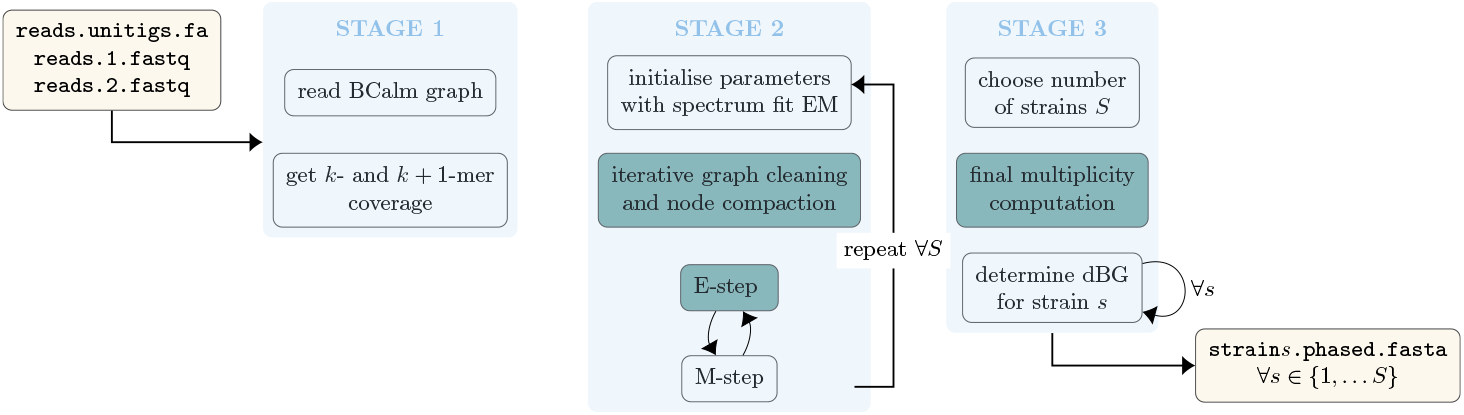
Overview of the HaploDetox pipeline, dark green highlights denote where the CRF is used.

### Conditional Random Field

A Conditional Random Field (CRF) is a Probabilistic Graphical Model that encodes the following joint conditional probability [11]:

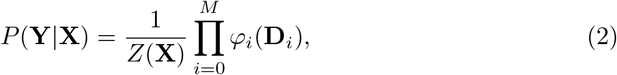

with **Y** a set of unknown variables, **X** a set of observed variables and **D**_*i*_ ⊆ **X** ∪ **Y** such that each **D**_*i*_ contains at least one element from **Y**. The *factors φ*_*i*_ express probabilistic relations between the variables in their *scope* **D**_*i*_. The factors are not required to express probability distributions (i.e. the values they assign to the range of possible values of *D*_*i*_ do not have to sum to one). However, the complete product is normalised by the *partition function* 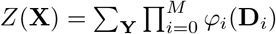 .

We define our CRF for a fixed number of strains *S* (see Fig. 3 for a visualisation). There is an unknown strain-specific multiplicity variable 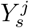 for each node and arc *j* and *s* ∈ {1, …, *S*}. Additionally, there is an observed variable *X*^*j*^ for each node and arc *j* that expresses the (average) coverage of that node/arc. We then define *coverage factors* 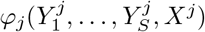 that quantify the relationship between observed coverage of the nodes and arcs *j* and their likely multiplicities *m* = (*m*_1_, …, *m*_*S*_) as follows:

**Fig. 3.**
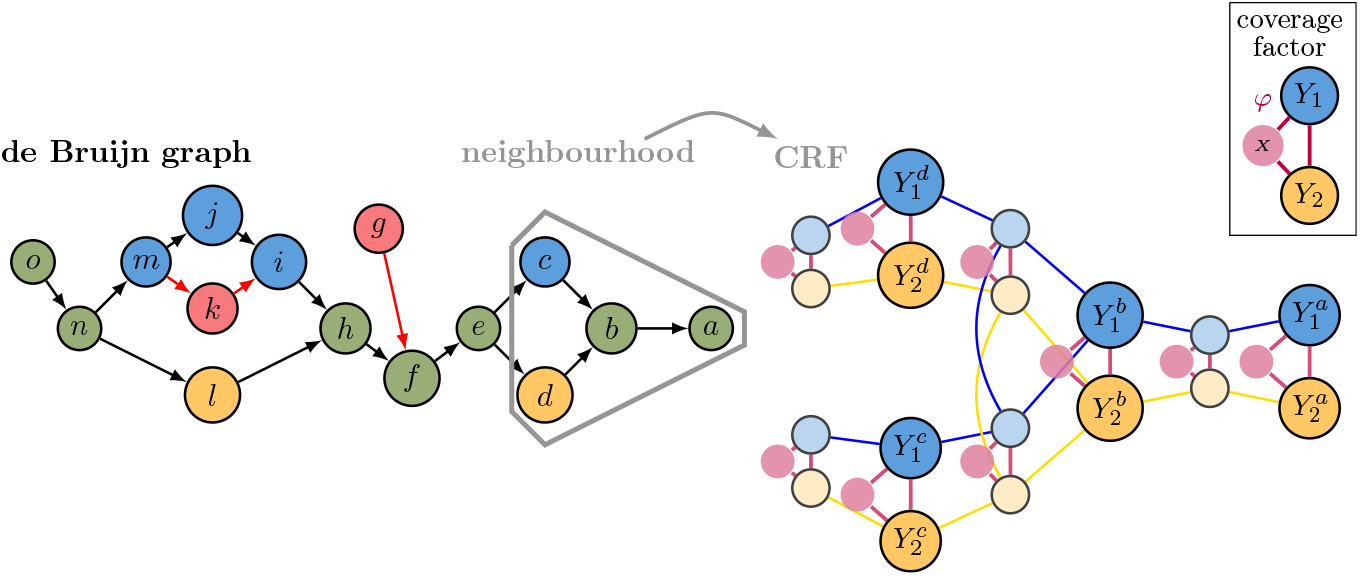
Small example de Bruijn graph with a neighbourhood of size 1 selected around node *b*. For this neighbourhood the corresponding Conditional Random Field is shown under the assumption that the data contains two different strains. The CRF contains unobserved multiplicity variables 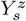 for each node/arc *z* and for each assumed strain *s* and observed variables *X* ^*z*^ for each node/arc. cliques formed by 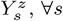, ∀*s* and *X*^*z*^ arise from the coverage factors (purple connections), while the connections between variables corresponding to different nodes/arcs in the graph arise because conservation of flow of multiplicity is enforced in all strains (blue connections for strain 1 and yellow connections for strain 2)

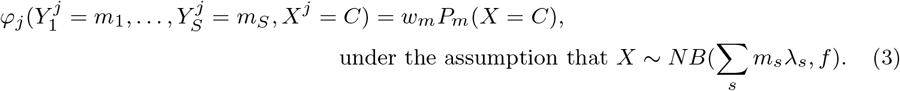

These coverage factors rely on a mixture of negative binomial distributions fitted to the observed *k*-mer coverage spectrum (see Fig. 1(b)). We assume that each strain *s* has an average coverage *λ*_*s*_ = *π*_*s*_*λ* such that ∑ _*s*_ *π*_*s*_ = 1 and *λ* is the average coverage estimate for those *k*-mers that occur exactly once in each strain genome. Theoretically, the number of sequencing reads that cover a certain position in the genome is a Poisson random variable and the *k*-mer coverage for each **m** should follow a Poisson distribution with mean ∑ _*s*_ *m*_*s*_*λ*_*s*_. However, sequencing biases lead to a variance that is larger than the mean. Therefore we model the coverage given a certain multiplicity *m* with a negative binomial distribution with mean *µ* = ∑ _*s*_ *m*_*s*_*λ*_*s*_ and overdispersion factor *f* such that the variance is equal to *fµ*. The negative binomial distribution representing the erroneous *k*-mers (multiplicity (0, …, 0)) is fitted with a separate mean *µ*_0_ and overdispersion *f*_0_. Because we assume that the strains in the sample are highly similar, we fit the model such that the highest peak in the *k*-mer spectrum corresponds with these unique *k*-mers shared by all strains. Additionally, we assume that all strains are equally diverged such that peaks arising from *k*-mers unique to a strain are of similar height (see Fig. 1(b)). The mixture model is trained iteratively using an EM-procedure that is described fully in Additional file 1, Appendix A.1. Finally we define *conservation of flow factors* for all nodes *n* and all strains *s*:

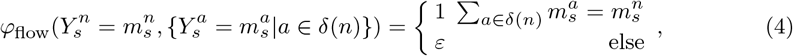

with *δ*(*n*) the set of arcs incoming to or outgoing from node *n* and *ε* ≪ 1.

### Graph cleaning

For our graph cleaning step, we imply conservation of flow at the nodes of the (unsimplified) de Bruijn graph while we simultaneously account for possible erroneous nodes caused by sequencing errors. In Steyaert et al. [12] we showed that using contextual information in the form of conservation of flow improves error detection and it is thus valuable to imply this property already during graph cleaning. We, therefore, use a CRF on a subgraph around nodes/arcs that are potentially erroneous to determine if they should be removed from the graph. By using a single model to detect strains and sequencing errors simultaneously, we avoid the bias and errors introduced when performing error-removal and strain detection separately.

### Determining the number of strains

We run stage 2 of the HaploDetox pipeline (see Fig. 2) for different assumptions of the number of strains *S* present in the dataset. At the end of stage 2 we compute several model fit measures (see Additional file 1, Appendix A.2, Table A1). We choose the model with the lowest BIC-score of the three models (see also a comparison of performance of model fit measures in Additional file 1, Appendix A.2). Because stage 2 contains a graph cleaning procedure, the number of elements based on which the models for different strain numbers are fitted will differ. The likelihoods that can be obtained by approximate inference on the CRF would therefore not provide good model selection criteria. On the other hand, we do want to have cleaned the graph based on the most likely number of strains. We attempted model selection based on the initialisation model on all nodes before graph correction, but this did not provide good results. For this reason we evaluate the model as follows: we view the obtained multiplicity assignments as a model-based clustering into multiplicity bins, which we evaluate based on a likelihood computed only over those variables corresponding to nodes that occur in all graphs. The likelihood considers only the fitted mixture model:

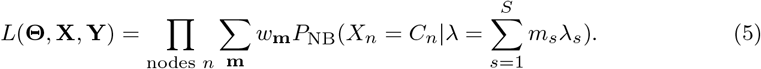

### Strain-specific de Bruijn graphs

After a final multiplicity computation for all nodes and arcs using the model for the chosen number of strains, we extract a de Bruijn graph for each assumed strain. Based on the initial de Bruijn graph we obtain the de Bruijn graph for strain *s* as follows: we remove all nodes that have an estimated *m*_*s*_ = 0 and concatenate the thus created new linear, unbranching paths in the graph. These separated graphs could be used to obtain an assembly, but for now we do not attempt any repeat resolution. We output each unitig (node) in the graph as a (relatively small) contig.

### Comparison to other methods

We compare our method to EVORhA [6], BHap [7], and mixtureS [8]. These three tools are the most similar to HaploDetox in terms of input data, (i.e., a short read dataset comprising multiple strains of a single species) and scope (i.e., whole genome haplotype reconstruction). All three tools present their reconstruction in the form of variant calls with respect to a reference genome of the species in the sample. To obtain such results based on the contig output from HaploDetox, we use Minimap2 [15] to align the contigs from the phased graphs to the reference genome and paftools from the Minimap2 package to call variant positions. We only take into account SNPs because BHap and mixtureS are limited to SNPs.

### Datasets

The first type of datasets are constructed as follows: we start from a single reference genome fasta file. Given the desired number of strains (2, 3 or 4) and a total number of SNPs divided over all strain genomes (100, 200, 500, 1000, 2000, 5000), a random phylogenetic tree is sampled based on which the number of SNPs is divided over the strains. The root of the tree never contains SNPs, but if *n*_strains_ ≥ 3, two or more strains can share SNPs. SNPs are inserted at random positions in the reference fasta file to obtain simulated strain genomes. From these simulated strain genomes reads are sampled using ART [16]. Finally, the simulated reads are combined into mixed datasets in which each strain is present at a different fraction of the total read support. We adopted the fractions per number of strains from [17], but removed those combinations where two strains received the same fraction, and we simulated 3 total sequencing depths (100×, 200× and 300×). Each combination of parameters was simulated for three different repetitions. These datasets are overly simplified compared to real-life datasets. However, all SNP positions that must be identified are known exactly. This allows for the most optimal characterisation of true-and false positive and negative results.

Secondly, we retrieved real Illumina read datasets from isolated strains of three different organisms that showed an almost perfect concordance with an existing finished RefSeq assembly (Table 1). These real reads were subsampled and mixed at differing proportions to obtain in silico mixed datasets. We use the same fractions as for the SNP simulated datasets, and we obtain 3 different total sequencing depths for all fractions. However, the maximum total sequencing depth depended on the sequencing depth of the isolate datasets. We additionally simulated reads using ART based on the reference genomes for the isolates, which are mixed at the same proportions as the read mixes. For the simulated datasets we are dependent on an alignment of the Ref-Seq assemblies to a chosen reference genome to determine the ground truth variants to be detected. To this end we have used Mummer’s dnadiff function [18]. For the real read mixed datasets, we obtain variant calls based on the original single genome read sets using Minimap2 and paftools [15].

**Table 1.** Datasets used to create in silico mixtures and their accession numbers.

| species | strain | read accession no. | reference accession no. | used as |
| --- | --- | --- | --- | --- |
| *M. tuberculosis* | GG-5-10 | SRR6389903 | GCF_002886165.1 | strain 1 |
|  | GG-109-10 | SRR6389909 | GCF_002887255.1 | strain 2 |
|  | GG-45-11 | SRR6389911 | GCF_002887145.1 | strain 3 |
|  | GG-111-10 | SRR6389902 | GCF_002886145.1 | strain 4 |
|  | H37Rv |  | GCF_000195955.2 | reference |
| *P. aeruginosa* | FDAARGOS 610 | SRR9160467 | GCF_006364795.1 | strain 1 |
|  | FDAARGOS 501 | SRR8178086 | GCF_003812905.1 | strain 2 |
|  | FDAARGOS 767 | SRR9641503 | GCF_006364735.1 | strain 3 |
|  | FDAARGOS 571 | SRR8179109 | GCF_003812885.1 | strain 4 |
|  | PAO1 |  | GCF_000006765.1 | reference |
| *S. enterica* | Bareilly str. CFSAN000752 | SRR5972872 | GCF_004848685.1 | strain 1 |
|  | Typhi str. 2018K-0756 | SRR10279247 | GCA_008705215.1 | strain 2 |
|  | Heidelberg str. 11-004736-1-7 | SRR3707412 | GCF_016452005.1 | strain 3 |
|  | Newport str. SAP18-8729 | SRR8548649 | GCF_004224885.1 | strain 4 |
|  | Typhimurium str. LT2 |  | GCF_014334155.1 | reference |

Finally, we retrieved 5 different datasets from Sobkowiak et al. [19] that contain short reads of in vitro mixed clinical *M. tuberculosis* isolates (proportions 0.95-0.05, 0.9-0.1, 0.7-0.3, 0.3-0.7, 0.1-0.9, and 0.05-0.95). The isolates were sequenced as well. Thus, even though we lack an assembled strain genome as ground truth, we can use the reads of the isolates to obtain ground truth SNP calls. These datasets give the best representation of a real mixed dataset we could find that still provides a characterised reference. The datasets are available on the European Nucleotide Archive (ENA) under project PRJEB2794, see Additional file 1, Appendix C for the accession numbers of each dataset.

### Evaluation metrics

In the datasets for which we have an assembled ground truth genome available, we determine for each node and arc its true multiplicity *m*_GT_ by aligning all ground truth strain genomes to the de Bruijn graph. Based on this, we call a multiplicity assignment *m* correct whenever the correct strains are present in the assignment (i.e. 𝟙 (*m*_*s*,GT_ *>* 0) = 𝟙 (*m*_*s*_ *>* 0) ∀*s* = 1 … *S*). We calculate a strain assignment accuracy as

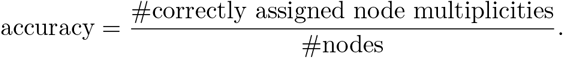

We calculate the strain assignment accuracy based only on those nodes with multiplicity **m** = (*m*_1_, …, *m*_*s*_) such that ∃*s, m*_*s*_ = 0 and ∃*s*^′^, *m*_*s′*_ ≠ 0. In other words, we do not take the erroneous nodes nor the nodes present in each strain into account when computing the accuracy. These two classes contain the majority of the nodes and are more easily classified correctly. Including such nodes results in very high accuracies that are less easily compared.

SNP calls are evaluated using precision, recall, and F_1_ metrics based on the evaluation methodology of [5]. Because the estimated number of strains might differ from the true number of strains, we determine which predicted strain shows the highest similarity to a true strain. To this end, we calculate jaccard scores for an all to all comparison of the predicted strains and the ground truth strains. We determine the most similar true strain for each predicted strain (as evaluated by the jaccard score). If more than one predicted strain is assigned to a certain true strain, we only keep the prediction with the highest jaccard score. We compute *TP* as those SNPs predicted in a strain that are present in the ground truth (GT) strain, *FP* as those SNPs predicted in a strain that are not present in the GT strain and *FN* as those SNPs that are present in the GT strain, but were not predicted in the strain estimate.

Finally, we evaluate the model fit of the mixture models used by HaploDetox and the methods in our comparison. On the one hand, we check whether the estimated number of strains is equal to the true number of strains. On the other hand, we evaluate the estimated strain abundances using the mean absolute error (MAE) between the true fractions (*π*_*s*_) and the estimated fractions 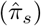, as in [5, 6, 8]:

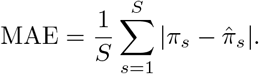

We compute the MAE based only on the most similar estimated strain for each true strain, if a true strain had no most similar estimated strain assigned we set 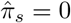.

## Results

We evaluate HaploDetox in terms of node multiplicity assignment accuracy on its own and based on strain-aware SNP calls in comparison with existing methods. We first explore performance on simulated read data based on simulated strains. Then we look at performance on several real read in silico mixtures for which we also created corresponding simulated datasets. Finally, we look at an in vitro mixture dataset, as this provides the closest resemblance to a real-world situation, while still providing a ground truth.

### Simulated SNPs, simulated reads

These datasets are the simplest presented here: the variation between strains consists solely of SNPs and the reference genome we use for genome alignment is the direct ancestor of all the strains: i.e. there is no part of the genome that might be present in all strains, but can still be called as a variant with respect to the reference sequence used for variant calling.

### Node assignment accuracy

Because we know the ground truth exactly in these simulated SNP datasets, we can evaluate the accuracy with which nodes are assigned to a certain multiplicity bin **m** = (*m*_1_, …, *m*_*S*_), when the assumed number of strains *s* is correct. We rerun the last stage of HaploDetox using the estimated model corresponding with the true number of strains even though the estimated number of strains might have been different in the model selection step. Fig. 4 shows the results of this evaluation, separated by number of simulated SNPs, but averaged over the different abundance combinations. The accuracy goes down as the number of strains increases (Fig. 4(a)). This is probably due to an increased complexity of the model that must be fitted. When there are more SNPs different between the strains, strain assignment accuracy increases. More SNPs allow for an easier detection of the peaks belonging to the individual strains and thus an easier model fit. Whenever there is a deviation from this trend in Fig. 4(a) (e.g. the box plot for the 3 strain, 2000 SNP simulation), we believe this to be caused by a highly uneven distribution of SNPs between the strains. The differences in difficulty of the model fit between simulation parameter settings is also reflected in the accuracy of the number of strains determination (Fig. 4(b)): the correct detection of the 4 strains model is challenging, even for the datasets with 2000 and 5000 simulated SNPs, the correct number of strains is selected in approximately 25% of the cases. The outliers in the boxplots (Fig. 4(a)) often represent datasets where two strain abundances are (almost) equal, or datasets where one of the abundances is very small. In the first case, two strains are easily confused with one another based on the coverage information, while in the latter case, the low abundance strain might be confused with sequencing errors. Additional file 1, Appendix B, Supplementary Fig. B2 also illustrates that accuracy is lowest when two strain fractions are close to one another. Finally, a higher total coverage leads to an increase in accuracy.

**Fig. 4.**
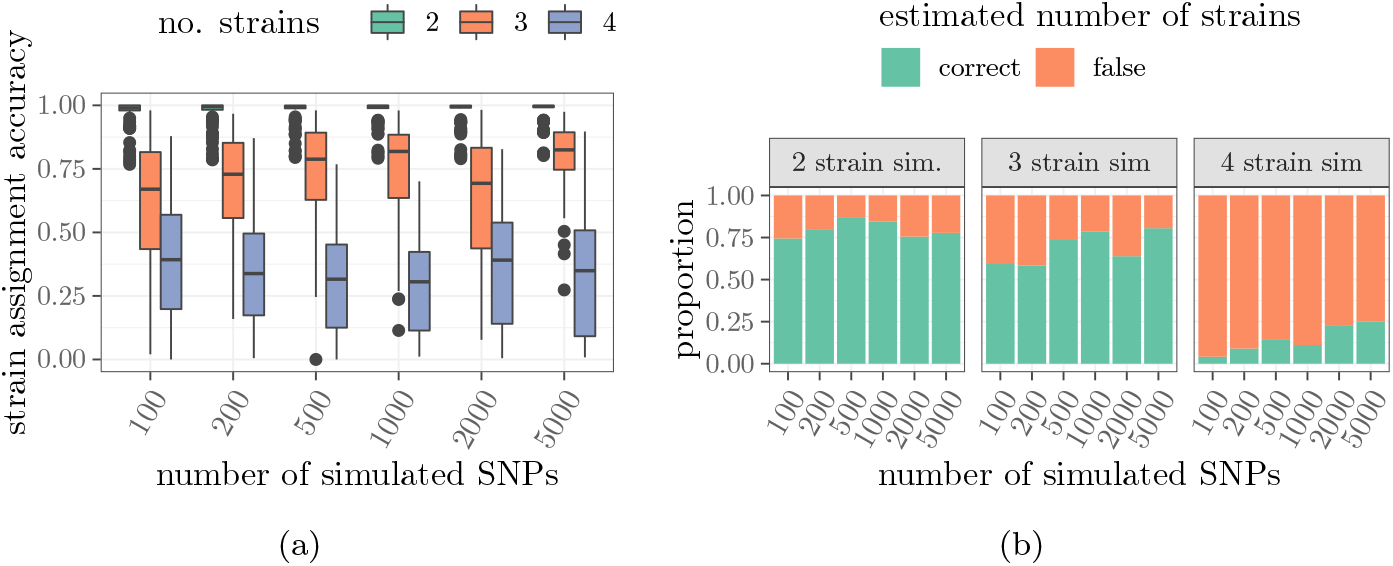
Accuracy evaluation for the simulated SNP datasets. **(a)** Strain multiplicity assignment accuracy. **(b)** Strain number estimation accuracy. For (a) only those nodes not belonging to all strains (shared region) or to no strains (sequencing errors) were considered. Results are averages (a) or counts (b) over several strain abundance combinations.

### Comparison with existing methods

HaploDetox reaches a higher F_1_ score than the best method in our comparison, mixtureS (Fig. 5). On the datasets with only two strains, HaploDetox always outperforms mixtureS both in terms of Precision and Recall. For the 3 and 4 strain datasets, we see that mixtureS has a higher precision on average than HaploDetox: HaploDetox is more prone to underestimating the number of strains, in which case it might collapse two strains, resulting in more false positive SNP calls that belong to a different strain that was not detected by HaploDetox. However, because of the higher recall of HaploDetox, we still obtain a higher F_1_ score average than mixtureS. BHap and EVORhA often overestimate the number of strains. We only count true positive SNP calls of the predicted strain with the closest resemblance to one of the ground truth strains. This results in a low recall when the strain number estimate is too high, but precision can still be high. EVORhA progressively merges clusters (each cluster corresponds to a potential strain). We believe this is done too conservatively, leading to one true strain being spread over several predicted strains by EVORhA. This leads to a disadvantage for EVORhA in the way we determine TP calls.

**Fig. 5.**
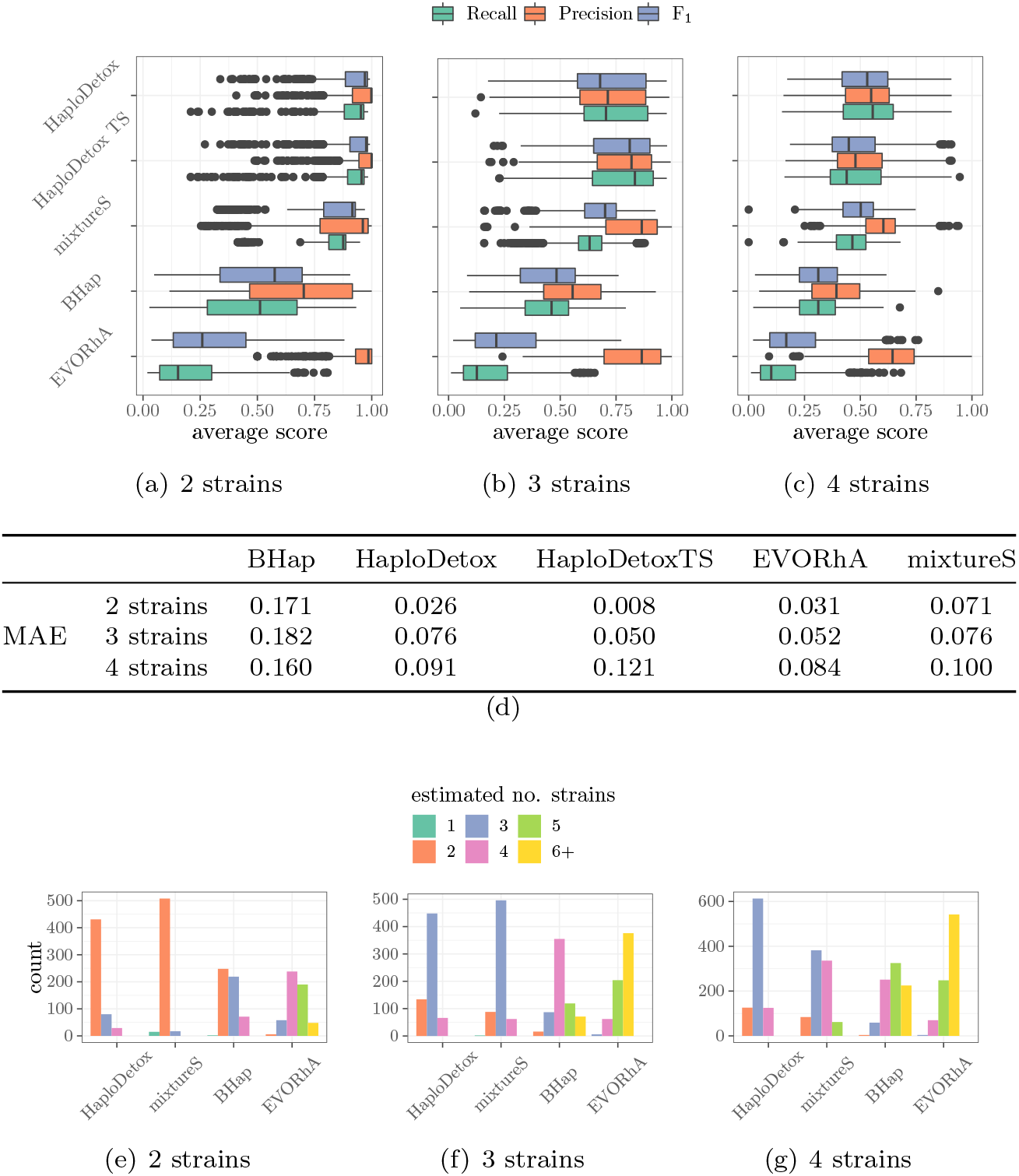
(a-c) Comparison with existing methods on the simulated SNP datasets in terms of F_1_-score, precision, and recall for 2 strains **(a)**, 3 strains **(b)**, and 4 strains **(c)** simulation. **(d)** Comparison in terms of average mean absolute error (MAE) per assumed number of strains, for each method tested. This is a measure for the accuracy of the strain fraction estimates, a lower value is better. **(e-g)** Comparison in terms of estimated number of strains for 2 strains **(e)**, 3 strains **(f)**, and 4 strains **(g)** simulation. Averages or totals are always taken over all different number of SNP and different fraction simulations. TS denotes that we passed the true number of strains to HaploDetox.

### In silico mixture

To the best of our knowledge there is no strain mixture dataset with more than two strains available that has its strains characterised independently such as the in vitro 2 strain mixture presented in the next section. Therefore, we use in silico mixtures based on real isolate genome bacterial reads to evaluate the performance of HaploDetox on real read mixes of 2, 3 and 4 strains. These datasets are more complex than the simulated SNP datasets because they contain other types of genomic variation between strains than only SNPs, and because real reads are often still harder to characterise than their simulated counterparts. Based on the assembled genomes of each single genome read set, we created a corresponding simulated read dataset for each simulated mixture at each simulated abundance. The results based on simulated data are given in Additional file 1, Appendix B, Supplementary Figures B3 and B4.

For real datasets, HaploDetox performs well when only 2 strains are present, but its strain assignment accuracy declines when the datasets contain 3 or 4 strains (Fig. 6). While we saw this trend in the simulated SNP datasets as well, it is more pronounced here. Additionally, *M. tuberculosis*, which has a higher repetitiveness, but also less variation between strains in the dataset seems to be the hardest to characterise. When we compare the results on the real reads (Fig. 6(a)) with those on the simulated read mixes of the reference assemblies (Additional file 1, Appendix B, Supplementary Fig. B4) we conclude that some of the reduced accuracy can be attributed to an imperfect correspondence between read set and assembly, especially in *M. tuberculosis*. However, the very low accuracy in the 4 strain datasets remains an issue.

**Fig. 6.**
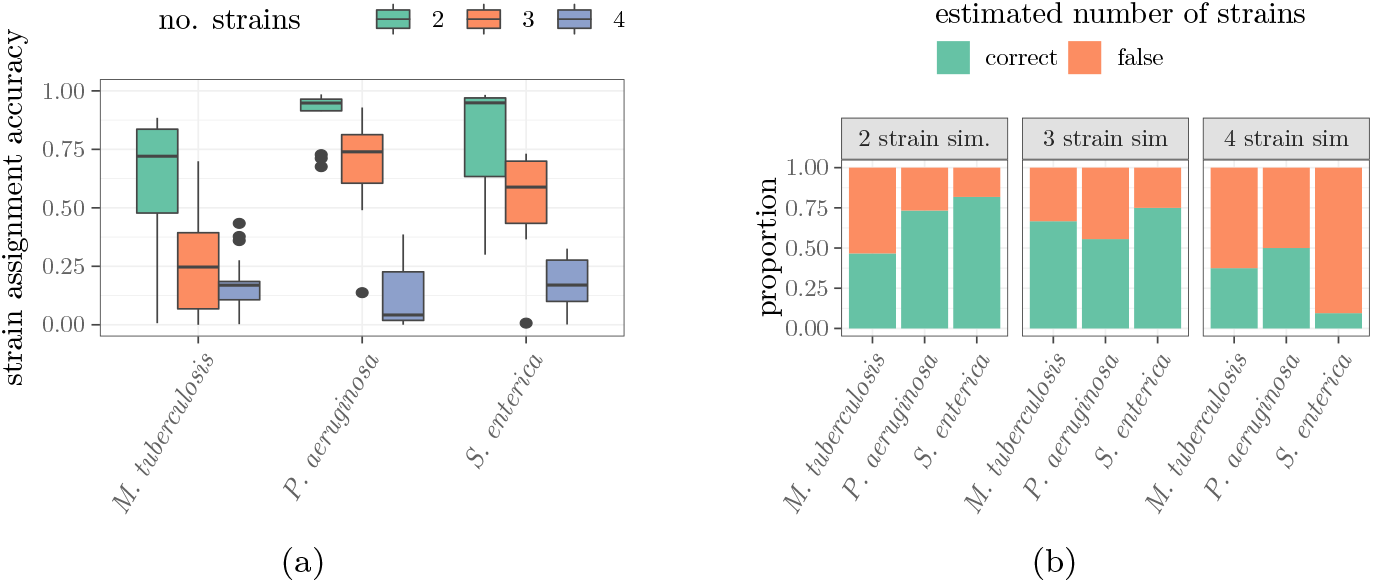
Accuracy evaluation for the in silico read mixes. **(a)** Strain multiplicity assignment accuracy. **(b)** Strain number estimation accuracy. For (a) those nodes belonging to all strains (shared region) or to no strains (sequencing errors) were not taken into account.

In the comparison with the other methods (Fig. 7), HaploDetox performs better than the other methods in detecting the correct underlying strain mixture model: the average MAE is lowest even when estimating the number of strains and HaploDetox most frequently uses the correct strain estimate. While mixtureS performed well in number of strains detection on simulated data, on real data its model estimation performance goes down. HaploDetox performs better than all existing methods in terms of F_1_ score on the 2 and 3 strains mixtures, on the 4 strain mixture mixtureS performs slightly better. However, as previously, we see a sharp decline in accuracy of all methods when the number of strains present in the datasets grows. We observe that our largest gains in recall over mixtureS are made because we are better in the detection of SNPs shared among all strains but different from the assumed wild-type allele in the reference genome used for mapping and variant calling. When we remove such SNPs from the ground truth we see that HaploDetox and mixtureS perform similarly (avg. F_1_ for HaploDetox: 0.832 (2 strains), 0.491 (3 strains), and 0.336 (4 strains), for mixtureS: 0.774 (2 strains), 0.458 (3 strains), and 0.331 (4 strains)). SNPs shared among all strains are not informative for strain separation, which might be the reason that mixtureS reports such SNPs so little. However, if we want to characterise complete genomes of the strains in the data, it is important to detect even the common SNPs where all strains differ from the reference genome.

**Fig. 7.**
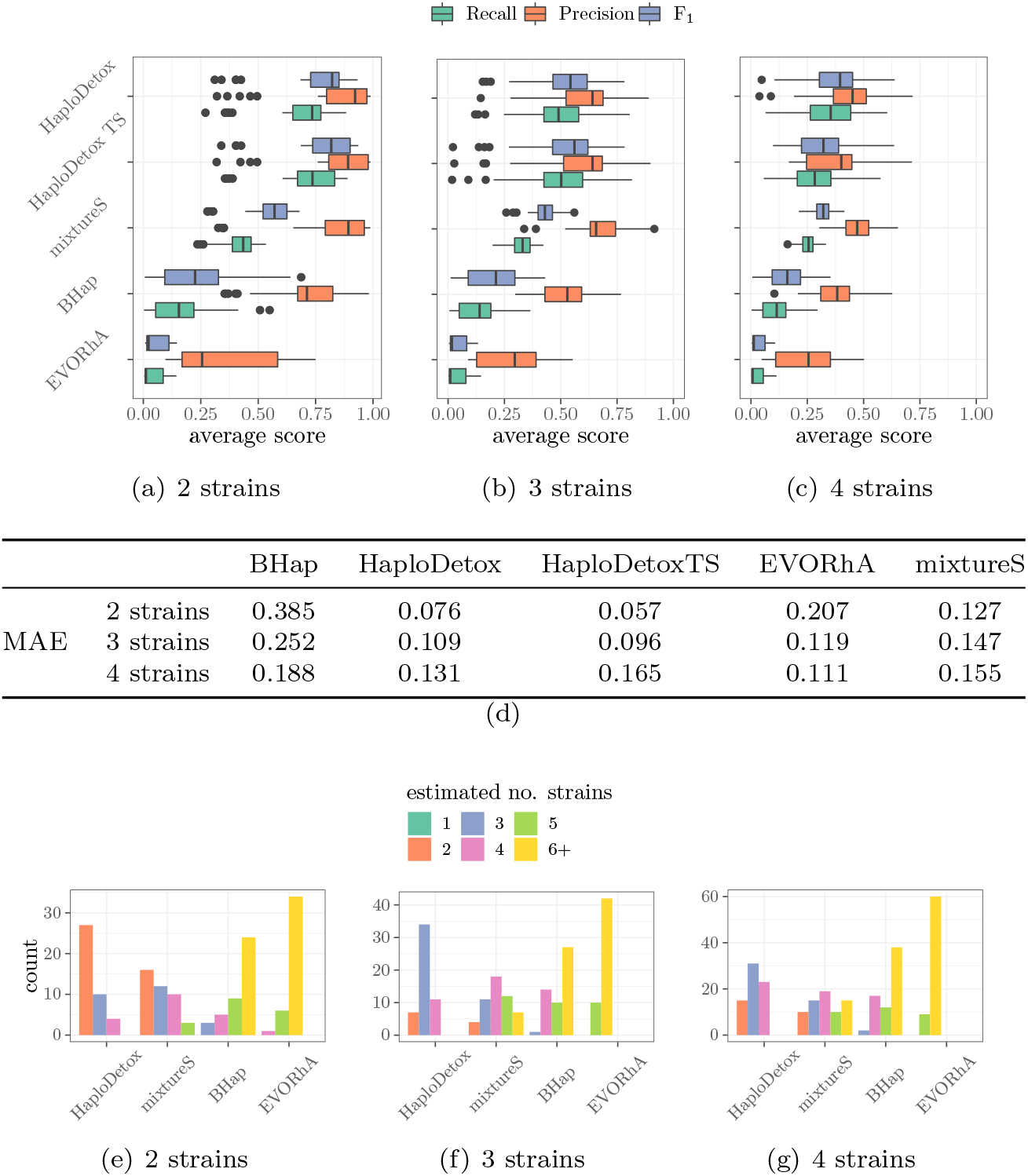
(a-c) Comparison with existing methods on in silico read mixes in terms of F_1_-score, precision, and recall for 2 strains **(a)**, 3 strains **(b)**, and 4 strains **(c)** simulation. **(d)** Comparison in terms of average mean absolute error (MAE) per assumed number of strains, for each method tested. This is a measure for the accuracy of the strain fraction estimates, a lower value is better. **(e-g)** Comparison in terms of estimated number of strains for 2 strains **(e)**, 3 strains **(f)**, and 4 strains **(g)** simulation. Averages or totals are always taken over all different organisms, total coverage, and different fraction simulations. TS denotes that we passed the true number of strains to HaploDetox.

### Runtime evaluation

The runtime of HaploDetox is influenced by several factors. First, a larger de Bruijn graph leads to a CRF with more variables and thus more computations necessary for one belief propagation iteration resulting in a larger runtime overall. E.g. in Table 2, *M. tuberculosis* has a much smaller graph than the other two organisms, resulting in lower runtimes. Second, the assumed number of strains for the model introduces a certain number of variables: 1 variable per assumed strain, per node/arc in the de Bruijn graph. For this reason we see that the runtime for the estimation of a model (almost) always increases with increasing number of strains assumption (Table 2). Finally, a bad model initialisation either because the number strains assumption is wrong, or because the initialisation EM converged to an erroneous initial estimate can influence the number of iterations needed by the approximate inference scheme and the number of iterations needed in the EM procedure. In Table 2, the runtime for *P. aeruginosa* for the 2 strains model is much longer when the true number of strains is 3 (erroneous model assumption) than when the true number of strains is 2 (correct model assumption) even though the number of EM iterations is on average the same and the number of nodes does not differ by the same factor as the increase in runtime. When the model assumption is wrong, approximate inference methods are much more likely to present oscillatory behaviour, resulting in slower convergence.

**Table 2.** Average runtime and number of EM iterations for the three different number of strain assumptions in the model selection stage. (TS: true number of strains, ES: assumed number of strains in model estimation)

| TS | ES | *M. tuberculosis* |  |  | *P. aeruginosa* |  |  | *S. enterica* |  |  |
| --- | --- | --- | --- | --- | --- | --- | --- | --- | --- | --- |
|  |  | nodes | iters | runtime | nodes | iters | runtime | nodes | iters | runtime |
| 2 | 2 |  | 6 | 0:15:43 |  | 4 | 0:36:30 |  | 3 | 0:32:52 |
|  | 3 | 22081 | 4 | 0:42:59 | 143308 | 5 | 2:31:42 | 113678 | 5 | 2:31:56 |
|  | 4 |  | 5 | 2:30:24 |  | 5 | 3:38:21 |  | 6 | 3:42:28 |
| 3 | 2 |  | 6 | 0:15:11 |  | 4 | 1:20:09 |  | 6 | 2:22:12 |
|  | 3 | 22135 | 5 | 0:54:51 | 177899 | 5 | 2:42:12 | 165545 | 6 | 3:03:01 |
|  | 4 |  | 4 | 2:20:46 |  | 4 | 3:09:54 |  | 5 | 3:16:59 |
| 4 | 2 |  | 5 | 0:13:38 |  | 4 | 1:15:17 |  | 7 | 3:06:33 |
|  | 3 | 26663 | 7 | 1:10:39 | 183863 | 5 | 2:54:54 | 200143 | 5 | 2:35:08 |
|  | 4 |  | 4 | 2:33:19 |  | 4 | 2:58:20 |  | 4 | 2:58:14 |

When we compare the runtime of HaploDetox with that of the other methods considered here (Table 3), we see that HaploDetox and mixtureS are much slower than BHap and EVORhA. Additionally, mixtureS is sometimes faster than HaploDetox, but at several instances, mixtureS’s model fit procedure suffers very bad convergence issues and its runtime becomes extremely high: mixtureS’s median runtime is between 1 and 2 hours, but the maximum runtime went up as high as 2 days, in contrast both HaploDetox’s mean and median runtime are around 5-6 hours (see also Additional file 1, Appendix B, Supplementary Fig. B5).

**Table 3.**
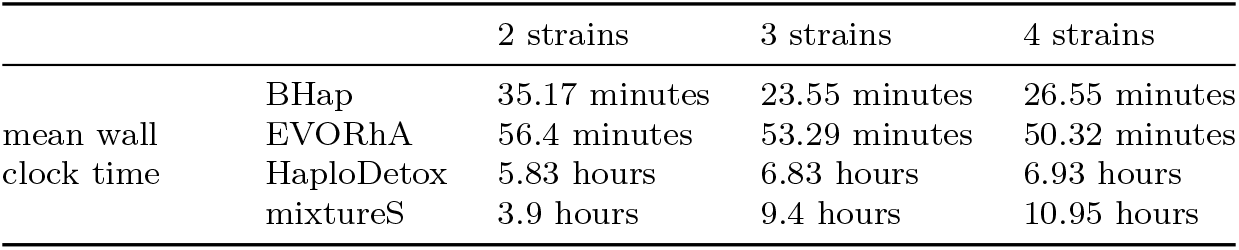
Comparison of HaploDetox’s total runtime with that of other methods.

### In vitro mixture

These final datasets [19] give the best representation of a real mixed dataset we could find that still provides a characterised reference. There is no reference assembly available, but we can determine ground truth SNPs based on the isolate read datasets. To detect both strains in the 0.05-0.95 and 0.1-0.9 datasets, we needed to change some default parameter settings. First, while our standard initialisation in HaploDetox is to evenly space the strain fractions, for this dataset we initialised the smallest strain fraction to a fixed value of 0.05 and evenly spaced the remaining strain fractions. This initialisation makes HaploDetox more sensitive to detecting low coverage strains, but might lead to fitting the lowest frequency distribution in the mixture to part of the erroneous nodes when all strains are highly covered. Second, we do not retrain the error distribution after graph cleaning, as this often leads to better accuracy when the number of variants between strains is low. However, the model selection procedure performs better when we retrain the error distribution together with the rest of the model after graph cleaning and this is therefore the default setting.

All methods overestimate the number of strains on many of the datasets (Fig. 8(b)). However, even when the number of strains is overestimated, HaploDetox performs best in terms of F_1_ score (Fig. 8(a)). Passing the true number of strains to HaploDetox reduces the variability in performance, but the median of the metrics remains similar. EVORhA has low recall and precision. On the previous datasets, we observed that EVORhA always has a low recall but often a precision similar to BHap. While mixtureS and BHap show a high precision, they suffer from a low recall. We noticed that the lower recall of mixtureS is reached because mixtureS almost never reports SNPs that are shared by all strains (Additional file 1, Appendix B, Supplementary Fig. B6). The reason for this might be that such SNPs are not informative for strain separation and that mixtureS therefore does not take them into account. Even these SNPs are, however, important if we aim to fully characterise a strain by its differences with a reference genome. Additionally, HaploDetox’s output can be extended to full haplotype-aware assembly, which gives it an additional advantage over mixtureS and BHap that are aimed solely at phasing SNPs.

**Fig. 8.**
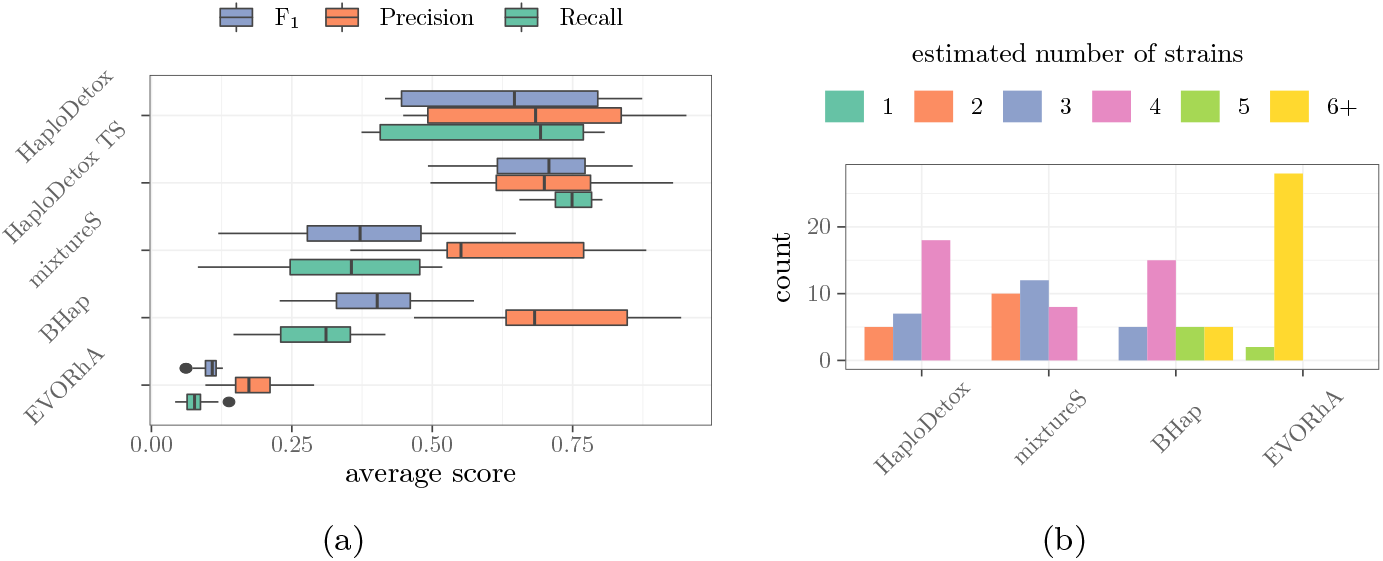
Comparison of HaploDetox to three other methods on the in vitro mixed datasets. Averages and totals are taken over 5 different in vitro mixed sets and 6 different abundance combinations (0.05-0.95, 0.1-0.9, 0.3-0.7, 0.7-0.3, 0.9-0.1, 0.95-0.05). TS denotes that we passed the true number of strains to HaploDetox. **(a)** Average F_1_, Precision and Recall. **(b)** Counts of the number of strains estimated for each dataset by the different methods.

## Discussion

We have shown that HaploDetox detects strain-specific SNPs with a similar or higher accuracy than existing methods. We observe that HaploDetox’s accuracy greatly depends on the model assumptions and the model initialisation. The standard parameter settings of HaploDetox were chosen such that our model selection performs best in terms of selecting the correct number of strains. However, different settings make our method more sensitive to low abundant strains or strains with a low divergence. For example, by default all distributions in the mixture model are retrained after graph cleaning. However, on the in vivo mixed dataset, we kept the error distribution from the initial model fixed while retraining the other distributions in the mixture model after graph cleaning. This improved SNP detection accuracy, but the number of strains was overestimated more often.

The performance of all methods goes down starkly as more strains are present in a dataset. At least for HaploDetox we believe that this is due to the increased complexity of the underlying *k*-mer spectrum model: for *s* strains and a maximum considered strain-specific multiplicity *M*, this model contains (*M* + 1)^*s*^ components. When the abundance of multiple strains is similar or when the sum of two strain abundances is similar to the abundance of a third strain, determining the most likely multiplicity based on the (*k*-mer) coverage becomes increasingly hard as the model contains many overlapping components. When none of the assignments have a high confidence in the correct value, even using flow to incorporate contextual information has less added value. Therefore, it could be beneficial to include other information that can phase strains in our method. By using a de Bruijn graph we lose a source of information: the fact that multiple variants might co-occur in a read or in the two mates of a pairedend read. While we initially motivated that, because of the lower mutation rate in bacterial strains, using read-overlap information is less valuable to phase reads than it is for viral haplotype reconstruction, it could still be valuable to research whether (paired-end) read information can be an additional source of information to phase haplotypes in the de Bruijn graph.

The runtime of HaploDetox is a limitation to the maximally detectable number of strains. Not only does the number of mixture components and thus possible values for the variables in the CRF increase with increasing number of strains assumed, the number of variables increases as well, because we create a variable for each assumed strain, for each node and arc in the de Bruijn graph. This leads to a substantial increase in necessary computations. To mitigate this, one could investigate whether it is possible to limit the number of multiplicity combinations considered for each node and arc in the graph. On the other hand, speeding up computations by for example looking into parallellised approximate inference techniques could also be part of the solution to our high runtimes. Finally, one could consider using an external method to get an estimate on the number of strains present in the dataset, such that HaploDetox only has to be run for one number of strains assumption. If this external method is highly accurate and the correct number of strains is detected, this would also avoid convergence difficulties due to bad model specification.

HaploDetox does not require a reference genome. However, it was developed to separate a small number of strains from a single species. Analysing a complete metagenomic dataset would require an alignment to a reference database of possible species present in the sample to bin reads per species. HaploDetox can then be run on the relevant bins to further separate the strains. When a reference database is not available, a (not necessarily strain-aware) metagenomic assembler can be used to obtain Metagenome Assembled Genomes (MAGs). Read bins obtained by mapping the reads to the MAGs can then be tested for the presence of multiple strains.

A potential extension of HaploDetox lies in using a combination of multiple time-point data or multiple otherwise related samples. Such multiple sample data is successfully exploited for de novo haplotype reconstruction by e.g. DESMAN [20], STRONG [21] and PoolHapX [22]. Also the authors of mixtureS recently showed that combining information of multiple pooled samples can improve strain phasing further [23].

Finally, we only evaluated HaploDetox’s performance in terms of strain-specific SNP calls. However, the general output of HaploDetox is a set of strain-specific de Bruijn graphs. This provides more options than just SNP detection: first, based on mapping and variant calling, all types of phased variants could be reported. Secondly, it would be interesting to implement repeat resolution and use the de Bruijn graphs to obtain a contiguous haplotype-aware assembly. Because HaploDetox’s maximally detected strain-specific multiplicity was now limited to *M* = 2, a first necessary step would be to detect the repeat multiplicities. After this, a repeat resolution method can be implemented to obtain full length contigs.

## Conclusion

We presented HaploDetox: a method that takes a set of second generation sequencing reads of a mixed-strain sample and returns a set of strain-specific de Bruijn graphs. We can detect strain-specific SNPs by mapping the unitigs of the strain-specific de Bruijn graphs to a reference genome for variant calling. For this SNP phasing task, we have a similar or even better performance than existing methods. Additionally, the output of HaploDetox is not limited to SNPs, but can be used to detect and phase all types of variants with respect to a reference. This is not possible with the best performing method in our comparison. Finally, to the best of our knowledge there is no method available to obtain a de novo haplotype-aware assembly of a bacterial mixed sample using short reads. We therefore believe the phased de Bruijn graphs are of value in the development of bacterial haplotype-aware assembly methods.

## List of abbreviations

BIC: Bayesian Information Criterion
CRF: Conditional Random Field
EM: Expectation-Maximisation
MAE: Mean Absolute Error
SNP: Single-Nucleotide Polymorphism

## Supplementary information

Additional information can be found in Additional file 1, containing Appendix A-C.

## Declarations

### Ethics approval and consent to participate

Not applicable

### Consent for publication

Not applicable

### Availability of data and materials

All data analysed during this study is previously generated, publicly available data. Accession numbers are provided in the main body of text.

### Availability and requirements

Our software code for HaploDetox is available with the following specifications:

Project name: HaploDetox

Project home page: https://github.com/biointec/haplodetox

Operating system(s): Linux

Programming language: C++11

Other requirements: cmake *>* 2.6.3, GCC *>* 4.7, Google’s sparsehash, LAPACK, Boost and GMP libraries

License: AGPL-3.0

### Authors’ contributions

AS and MVH contributed equally to this work. Conceptualisation by AS, KM and JF. Code implementation by MVH, AS, and JF. Analyses setup by AS, interpretation by MVH, AS and JF. Writing of the manuscript by AS and MVH. Project supervision by JF and KM. All authors read, edited and approved the manuscript.

## Acknowledgments

We thank the annonymous reviewers for their valueable feedback and suggestions on the manuscript.

## Authors’ information

Authors and Affiliations:

IDLab, Department of Information Technology, Ghent University-imec, Technologiepark-Zwijnaarde 126, 9052 Ghent, Belgium

Aranka Steyaert, Marie Van Hecke, Kathleen Marchal, Jan Fostier

Department of Plant Biotechnology and Bioinformatics, Ghent University, Technologiepark-Zwijnaarde 71, 9052 Ghent, Belgium

Marie Van Hecke, Kathleen Marchal

