## Supplementary material for "A Conditional Random Field approach for de novo reconstruction of bacterial haplotypes from a de Bruijn graph representation"

#### Appendix A Additional details on the HaploDetox pipeline

##### A.1 Training the negative binomial model

###### A.1.1 Initialisation

A good initialisation is essential to obtain good convergence properties of any Expectation Maximisation (EM) algorithm. We make several assumptions about the dataset to obtain an easy to compute, but accurate enough initialisation. The first assumption is that all strains share a large part of their genome such that the largest peak in the $k$-mer spectrum is the peak to which the distribution in the mixture with $m = (1, 1, \ldots, 1)$ should fit (see Fig. 1b in the main text). The second assumption is one of phylogenetic equidistance: all strains present will differ in approximately the same number of $k$-mers. These assumptions allow us to enforce equal weights for all peaks corresponding to the $k$-mers of a single strain. While this last assumption might not be true in practice when there are 3 or more strains present, we do observe that using this constraint enforces a quick and good convergence of the initial EM procedure. Given these assumptions, we initialise $\lambda_0 = 1.0$ and $\lambda$ to the average $k$-mer coverage and we run an EM procedure to fit a mixture of negative binomial distributions to the $k$-mer spectrum. This initialisation often gives a good enough fit to the $k$-mer spectrum, which we can use to remove erroneous nodes and arcs from the de Bruijn graph using the CRF to determine which nodes and arcs likely have a multiplicity $m = (0, \ldots, 0)$. After this removal, we finetune the model fit by estimating the multiplicities of the remaining nodes and arcs using the CRF and estimating the parameters in a second EM procedure based on these multiplicity estimates.

##### A.1.2 M-step estimation of mixture model

Because negative binomial maximum likelihood estimators do not have a closed-form solution, we use method of moments estimators in the M-step. Additionally, we add constraints based on assumed relationships between the different model parameters. We estimate a node and arc model separately, but the procedure is the same and we describe it only in terms of the nodes.

We start by separately fitting a truncated negative binomial distribution to the estimated error histogram. We denote the count of $k$-mers with estimated multiplicity $\mathbf{0} := (0, \ldots, 0)$, and (average) coverage $c$ as $\#_{\mathbf{0}}(c)$ which is calculated as:

$$\begin{aligned}\#_{\mathbf{0}}(c) \quad &= \sum_{\text{nodes } n} P(\text{mult}(n) = \mathbf{0})\,(\mathbb{1}(\lfloor \text{cov}(n) \rfloor = c)(1 - f_n) \\ &\qquad + \mathbb{1}(\lfloor \text{cov}(n) \rfloor + 1 = c)(f_n))\,,\end{aligned}$$

where $f_n = \text{cov}(n) - \lfloor \text{cov}(n) \rfloor$.

Next, denote the observed coverage of the nodes as $X = (X_1, \ldots, X_N)$, where $N$ is the number of nodes, and denote the expected multiplicity of a node $n$ as $\mathbf{m}^n = (m_1^n, \ldots, m_S^n)$, where $S$ is the assumed number of strains. We consider the estimation of the vector $\boldsymbol{\lambda} = (\lambda_1, \ldots, \lambda_S)$ such that the negative binomial mean for multiplicity $\mathbf{m}^n = (m_1^n, \ldots, m_S^n)$ is equal to $\sum_{s=1}^{S} m_s^n \lambda_s$. The vector $\boldsymbol{\lambda}$ can be estimated using an ordinary least squares (OLS) estimator to solve for $\hat{\boldsymbol{\lambda}} = \text{argmin}||\mathbf{M}\boldsymbol{\lambda} - X||^2$, with $\mathbf{M} = (\mathbf{m}^1, \ldots \mathbf{m}^N)$ an $N \times S$ matrix. The solution is thus obtained by solving the OLS problem $\hat{\boldsymbol{\lambda}} = ((\mathbf{M})^T\mathbf{M})^{-1}\mathbf{M}^T X$, which is done with a Cholesky decomposition.

After estimating $\boldsymbol{\lambda}$ for the node and arc coverage, we compute $\pi_1, \ldots, \pi_S$ such that $\lambda_s = \pi_s \lambda$ for both the node and arc model. We retain a single $\pi_1, \ldots, \pi_S$ such that $\pi_s = \frac{\pi_s(\text{node}) + \pi_s(\text{arc})}{2}$. While the actual coverage $\lambda_s$ might differ between nodes ($k$-mers) and arcs ($k + 1$-mers), the fractions should be the same in both models.

The overdispersion factor is assumed equal for all negative binomials in the mixture and is estimated using a weighted average of the component-wise estimates:

$$\hat{f} = \frac{1}{\sum_m w_m} \sum_m \sum_{\text{nodes } n} P(\text{mult}(n) = \mathbf{m}) \frac{(X_n - \sum_{s=1}^{S} m_s \hat{\lambda_s})^2}{\sum_{s=1}^{S} m_s \hat{\lambda_s}},$$

where $w_m = \sum_{\text{nodes } n} P(\text{mult}(n) = \mathbf{m})$, and the sum over $\mathbf{m}$ is a summation over all multiplicity combinations where $m_i \in [0, M]$. The maximum value considered is a parameter of the model and nodes with an estimated strain multiplicity $> M$ are not considered for parameter estimation.

To obtain the final weights for the mixture of negative binomials, we (optionally) imply several relationships between the weights, starting out from the matrix $w_m$. To this end we assume that $w_{m_i}$ and $w_{m_j}$ are equal when ordered$(m_i)$ = ordered$(m_j)$, using the assumption that all types of variants between strains are equally likely to occur in each strain. These constraints are combined with the weights calculated in

$w_m$ to obtain an overdetermined system that can be solved as a least squares problem. We use a LAPACK [1] routine to obtain a solution. While the equidistance assumption might not always hold in practice, we do observe that the added constraints maintain a more stable convergence to a reasonable solution.

#### A.2 Number of strains model selection

Table A1 provides an overview of the information criteria we considered (see also [2] for a description). Besides the well known AIC and BIC, we also compute several entropy based information criteria. The entropy of a model to the data is defined as:

$$E(n, K) = -\sum_{k=1}^{K}\sum_{i=1}^{n} t_{ik}\log t_{ik},$$

where $t_{ik}$ is the estimated posterior probability of element $i$ belonging to bin $k$. We tested the use of an entropy computed based on the mixture model alone, as well as an entropy where the $t_{ik}$ are determined by the CRF assignments. Note that almost all information criteria are minimised to determine the best model fit. The Entropy IC, however, is maximised.

**Table A1** Different information criteria considered for model selection. $d_K$ denotes the number of (independent) model parameters when assuming $K$ classes, while $n$ denotes the total number of elements to be classified and $E(n, K)$ is the entropy of a model with $K$ classes for $n$ elements. Note that the number of classes $K$ is not equal to the number of strains. The number of strains assumed will induce a certain number of distributions in the mixture model, each distribution represents a class. In HaploDetox we use the likelihood given by eq. (5) from the main text.

| | abbrev. | formula |
|---|---|---|
| Bayesian Information Criterion | BIC | $-2\log L(\mathbf{\Theta}_K, \mathbf{X}, \mathbf{Y}) + d_K \log(n)$ |
| Aikake's Information Criterion | AIC | $-2\log L(\mathbf{\Theta}_K, \mathbf{X}, \mathbf{Y}) + 2d_K$ |
| Integrated Complete Likelihood | ICL | $-2\log L(\mathbf{\Theta}_K, \mathbf{X}, \mathbf{Y}) + 2E(n, K) + d_K \log(n)$ |
| Classification Likelihood Information Criterion | CLC | $-2\log L(\mathbf{\Theta}_K, \mathbf{X}, \mathbf{Y}) + 2E(n, K)$ |
| Entropy Information Criterion | entropyIC | $1 - \frac{E(n, K)}{n\log(K)}$ |

Supplementary Fig. A1 shows the number of strains selection based on the different information criteria considered. In general, strain number selection is more often correct when there are more SNPs present between the different strains. When the true number of strains is 2, BIC and AIC tend to overestimate the number of strains slightly more often than the other criteria. However, when the true number of strains is 3 or 4, all other criteria underestimate the number of strains far more often. For this reason we chose to work with the BIC as criterion to select the strain number.

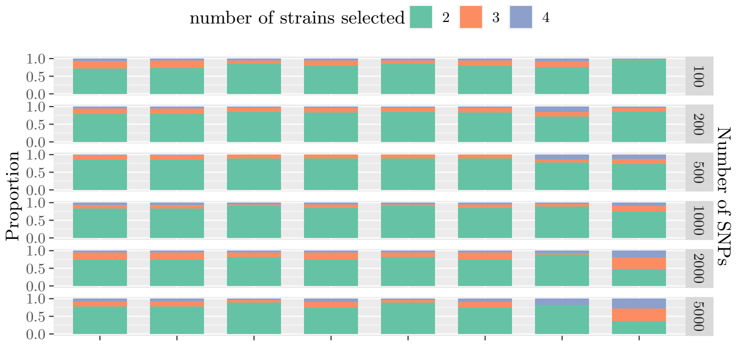


(a) 2 strain model selection outcomes

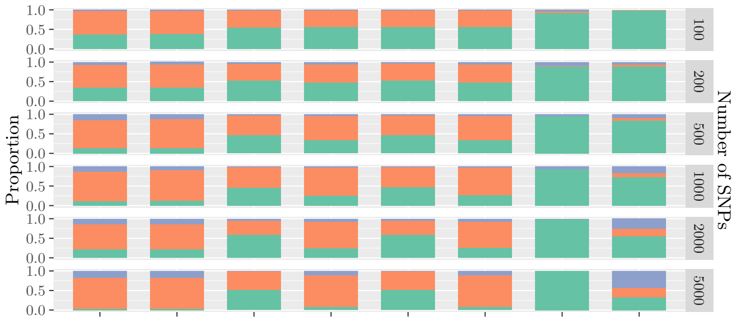


(b) 3 strain model selection outcomes

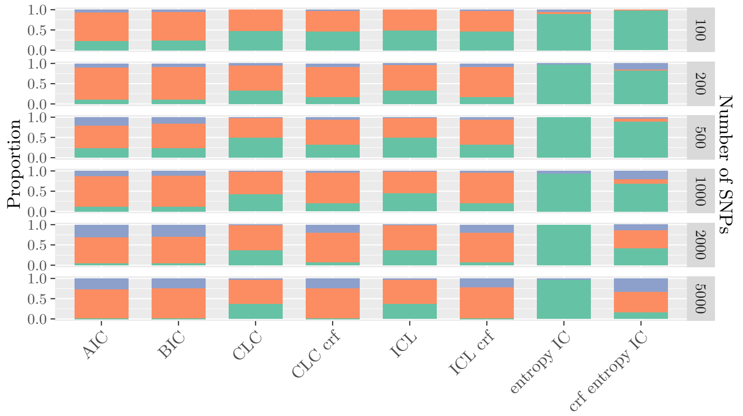


(c) 4 strain model selection outcomes

**Fig. A1** Evaluation of model selection criteria for the selection of number of strains present in the simulated SNP datasets. For CLC, ICL, and the entropy IC we considered entropy calculations based on the k-mer spectrum model alone or based on the CRF-assignments (crf).

### Appendix B Additional Figures illustrating results

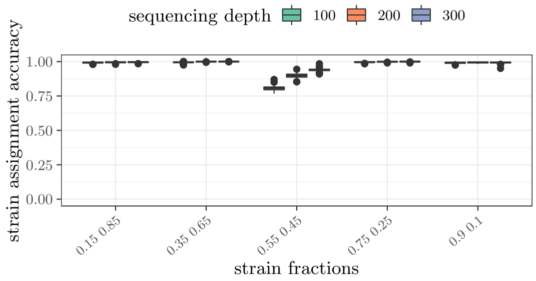


(a) 2 strains simulation

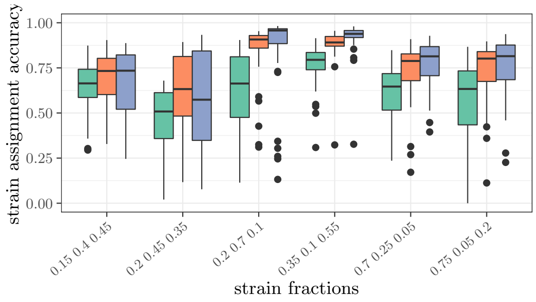


(b) 3 strains simulation

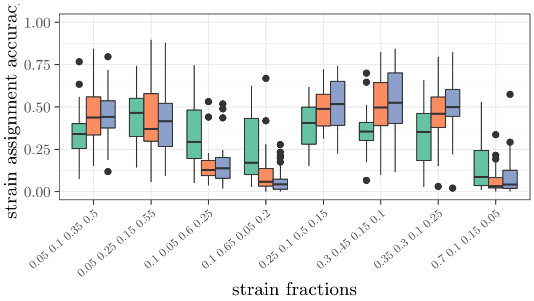


(c) 4 strains simulation

**Fig. B2** Strain assignment accuracy on simulated SNP datasets split by total simulated coverage (different coloured boxplots) and by simulated fraction of the total coverage for each of the strains (different sets of boxplots).

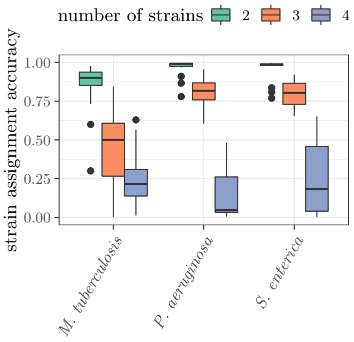


(a) Strain multiplicity assignment accuracy.

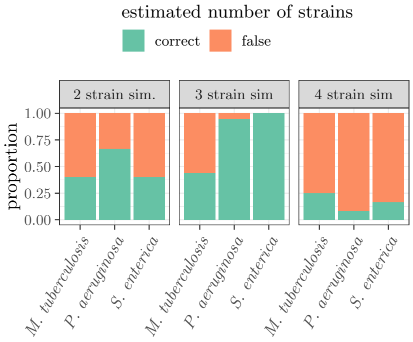


(b) Strain number estimation accuracy

**Fig. B3** Accuracy evaluation for the simulated read mixes. For (a) only those nodes not belonging to all strains (conserved region) or to no strains (sequencing errors) were considered.

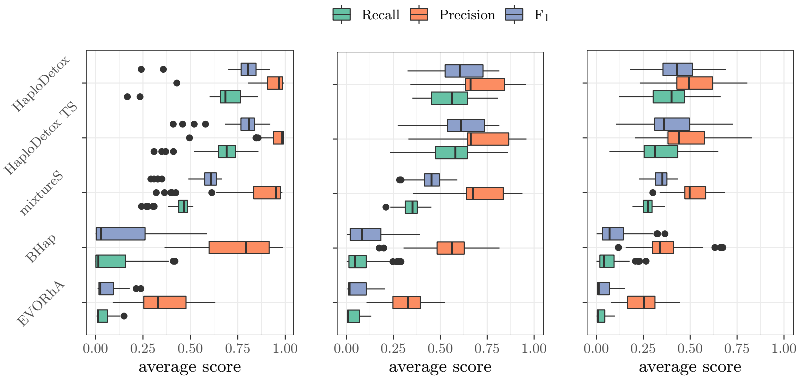


(a) 2 strains simulation (b) 3 strains simulation (c) 4 strains simulation

| | | BHap | HaploDetox | HaploDetoxTS | EVORhA | mixtureS |
|---|---|---|---|---|---|---|
| MAE | 2 strains | 0.243 | 0.078 | 0.016 | 0.226 | 0.058 |
| | 3 strains | 0.242 | 0.081 | 0.070 | 0.149 | 0.122 |
| | 4 strains | 0.231 | 0.101 | 0.121 | 0.139 | 0.136 |

(d) Average mean absolute error (MAE) per assumed number of strains, for each method tested. This is a measure for the accuracy of the strain fraction estimates, a lower value is better.

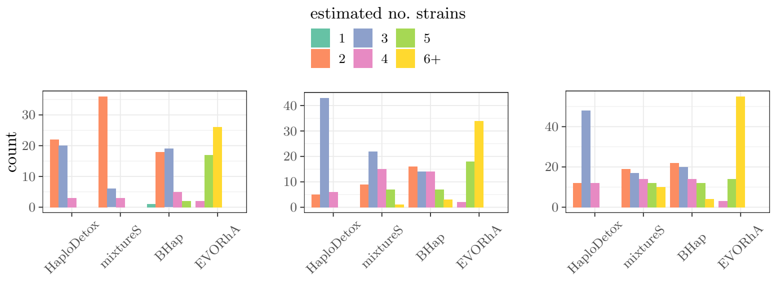


(e) 2 strains simulation (f) 3 strains simulation (g) 4 strains simulation

**Fig. B4** Comparison with existing methods on simulated read mixes in terms of $F_1$-score, precision, and recall (top), MAE (middle) and estimated number of strains (bottom). Averages or totals are always taken over all different organisms, total coverage, and different fraction simulations. TS denotes that we passed the true number of strains to HaploDetox.

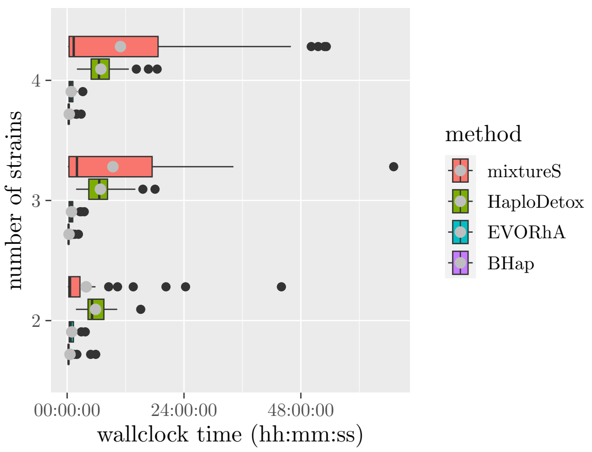


**Fig. B5** Comparison of total wallclock time used by all methods. Boxplots represent quantiles, grey dots represent mean.

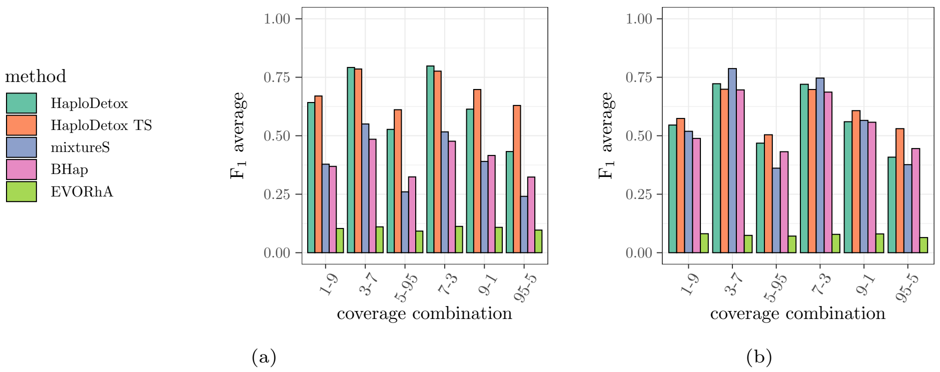


**Fig. B6** $F_1$ score per coverage combination, averaged over 5 in vitro mixed datasets. (a) $F_1$ computed with all SNPs of the strains with respect to a reference genome. (b) $F_1$ computed with the SNPs where both strains share the same mutation, but differ from the reference genome removed from the ground truth set.

### Appendix C Sobkowiak, et al. accession numbers

| | isolate 1 | isolate 2 | 0.05-0.95 | 0.1-0.9 | 0.3-0.7 | 0.7-0.3 | 0.9-0.1 | 0.95-0.05 |
|---|---|---|---|---|---|---|---|---|
| set 1 | ERR221642 | ERR221639 | ERR221638 | ERR221636 | ERR221643 | ERR221641 | ERR221640 | ERR221637 |
| set 2 | ERR221622 | ERR221624 | ERR221626 | ERR221625 | ERR221623 | ERR221627 | ERR221620 | ERR221621 |
| set 3 | ERR221661 | ERR221667 | ERR221666 | ERR221660 | ERR221663 | ERR221662 | ERR221665 | ERR221664 |
| set 4 | ERR221646 | ERR221648 | ERR221650 | ERR221649 | ERR221647 | ERR221651 | ERR221644 | ERR221645 |
| set 5 | ERR221657 | ERR221658 | ERR221655 | ERR221652 | ERR221656 | ERR221659 | ERR221654 | ERR221653 |

**Table C2** Accession number of the different datasets used from [3]. Each row corresponds to datasets from the same isolates.
